# Parallel power posterior analyses for fast computation of marginal likelihoods in phylogenetics

**DOI:** 10.1101/104422

**Authors:** Sebastian Höhna, Michael J. Landis, John P. Huelsenbeck

## Abstract

**Motivation:** In Bayesian phylogenetic inference, marginal likelihoods are estimated using either the path-sampling or stepping-stone-sampling algorithms. Both algorithms are computationally demanding because they require a series of power posterior Markov chain Monte Carlo (MCMC) simulations. Here we introduce a general parallelization strategy that distributes the power posterior MCMC simulations and the likelihood computations over available CPUs. Our parallelization strategy can easily be applied to any statistical model despite our primary focus on molecular substitution models in this study.

**Results:** Using two phylogenetic example datasets, we demonstrate that the runtime of the marginal likelihood estimation can be reduced significantly even if only two CPUs are available (an average performance increase of 1.96x). The performance increase is nearly linear with the number of available CPUs. We record a performance increase of 11.4x for cluster nodes with 16 CPUs, representing a substantial reduction to the runtime of marginal likelihood estimations. Hence, our parallelization strategy enables the estimation of marginal likelihoods to complete in a feasible amount of time which previously needed days, weeks or even months.

**Availability:** The methods described here are implemented in our open-source software `RevBayes` which is available from http://www.RevBayes.com.

**Supplementary information:** Supplementary data are available at *Bioinformatics* online.

## Introduction

Model selection in Bayesian phylogenetic inference is performed by computing Bayes factors, which are ratios of the marginal likelihoods as for two alternative models (Kass and Raftery, 1995; Sullivan and Joyce, 2005). The Bayes factor indicates support for a model when the ratio of the marginal likelihoods is greater than one. This procedure is very similar to likelihood ratio tests with the difference being that one averages the likelihood over all possible parameter values weighted by the prior probability rather than maximizing the likelihood with respect to the parameters (Posada and Crandall, 2001; Holder and Lewis, 2003). More specifically, the marginal likelihood of a model, *f*(*D|M*), is calculated as the product of the likelihood, *f*(*D|θ,M*), and the prior, *f*(*θ|M*), integrated (or marginalized) over all possible parameter combinations,

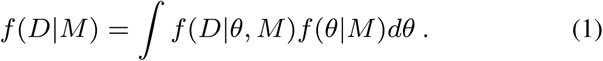

In the context of Bayesian phylogenetic inference, this quantity is computed by marginalizing over the entire parameter space, namely over all possible tree topologies, branch lengths, substitution model parameters and other model parameters (Huelsenbeck *et al*., 2001; Suchard *et al*., 2001).

The computation of the marginal likelihood is intrinsically difficult because the dimension-rich integral is impossible to compute analytically. Monte Carlo sampling methods have been proposed to circumvent the analytical computation of the marginal likelihood (Gelman and Meng, 1998; Neal, 2000). Lartillot and Philippe (2006) introduced a technique called thermodynamic integration, (also called path-sampling; Baele *et al*., 2012), to approximate the marginal likelihood. A similar method, stepping-stone-sampling (Xie *et al*., 2011; Fan *et al*., 2011), has more recently been proposed (see also Baele *et al*., 2012; Baele and Lemey, 2013; Friel *et al*., 2014, for a summary and comparison of these methods). The fundamental idea of path-sampling and stepping-stone-sampling is to use a set of *K* importance distributions, or power posterior distributions, from which likelihood samples are taken (Gelman and Meng, 1998; Neal, 2000; Lartillot and Philippe, 2006; Friel and Pettitt, 2008). The sampling procedure for each importance distribution is performed by a Markov chain Monte Carlo (MCMC) algorithm. That is, instead of running a single MCMC simulation as commonly done to estimate posterior probabilities (Huelsenbeck *et al*., 2001, 2002), *K* (usually between *K* = 30 and *K* = 200) MCMC simulations are needed to estimate the marginal likelihood of a model of interest. Obviously, this strategy can be very time consuming considering that a single MCMC simulation may take from hours to several weeks of computer time. The high computational time poses a major challenge for Bayes factor computations for many important problems, for example, comparing molecular substitution models (Posada and Crandall, 2001), selecting between complex diversification rate models (FitzJohn, 2012), and evaluating competing continuous trait processes (*e.g.*,Uyeda and Harmon, 2014).

In the present article we demonstrate how power posterior simulations can be performed on parallel computer architectures and report the achieved computational gain. The idea of parallel power posterior simulations is very similar to parallel Metropolis coupled MCMC algorithm (Altekar *et al*., 2004), with the important difference that power posterior simulations can be parallelized even more easily because no communication between processes is necessary. Additionally we show how our parallelization scheme can combined with existing parallelization techniques for distributed likelihood computation (*e.g.*, Aberer *et al*., 2014) to maximize usage of available CPUs.

## 2 Methods

The algorithm underlying path-sampling and stepping-stone-sampling can be separated into two steps: (1) likelihood samples are obtained from a set of K power posterior simulations; and (2) the marginal likelihood is approximated either by numerical integration of the likelihood samples over the powers (path-sampling) or by the likelihood ratio between powers (stepping-stone-sampling). The first step is the same for both methods and is the computationally expensive part. Thus, once samples from the power posterior distributions are obtained, it is possible to rapidly compute both the path-sampling and stepping-stone-sampling marginal likelihood estimates.

### 2.1 Power posterior sampling

Both, stepping-stone-sampling and path-sampling, construct and sample from a series of importance distributions. Lartillot and Philippe (2006) define the importance distributions as power posterior distributions, which are obtained by modifying the posterior probability density as

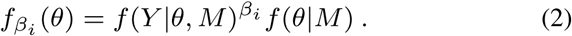

Here, *β* represent a vector of powers between 0 and 1. Then, for every value of *β*_*i*_ a draw from the power posterior distribution is needed and its likelihood score, *l*_*i*_, is recorded (Lartillot and Philippe, 2006; Friel and Pettitt, 2008). In principle, one such likelihood sample per power posterior distribution is sufficient, although multiple samples improve the accuracy of the estimated marginal likelihood considerably (Baele *et al*., 2012). We will use the notation *l*_*ij*_ to represent the *j*^*th*^ likelihood sample from the *i*^*th*^ power posterior distribution.

We illustrate the mean log-likelihood over different values of *β* in Figure 1. Commonly, the values of the powers *β* are set to the *i*^*th*^ quantile of a beta (0.3, 1.0) distribution (Xie *et al*., 2011; Baele *et al*., 2012). The rationale is that more narrowly spaced intervals are needed for the range of *β* where the expected likelihood changes most rapidly, *i.e.*, for *β* values close to 0 (Figure 1).

**Fig. 1.**
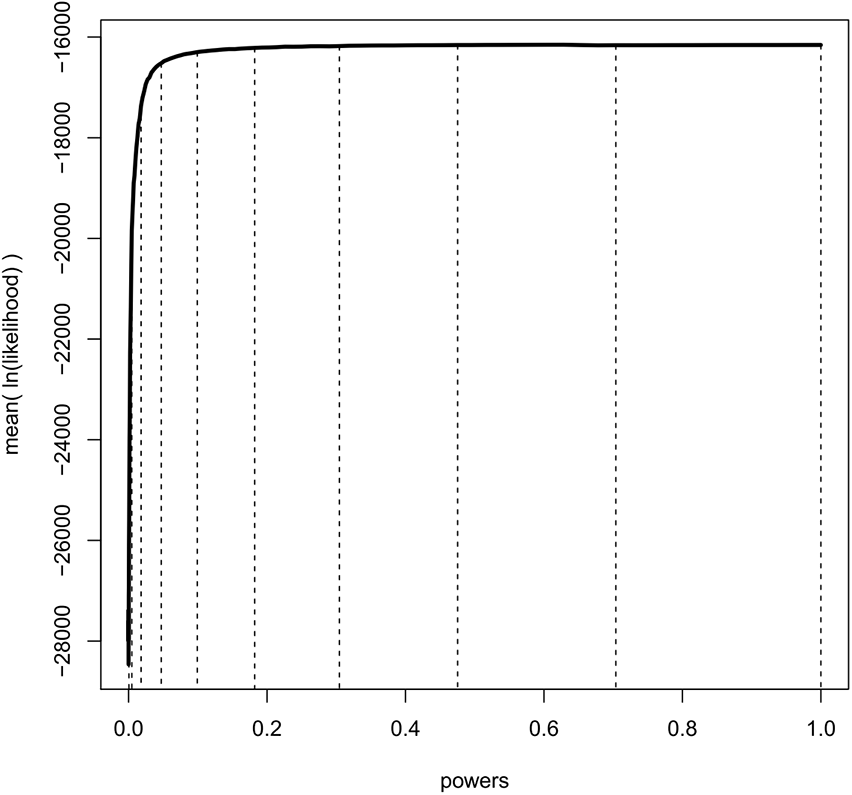
An example curve of mean log-likelihood samples over a range of different powers. The vertical, dashed lines show which values of powers were used when *K* = 11 and *β*_*i*_ = (*i*/(*K* − 1)) ^1.0/0.3^ for *i* ∈ {0, *K* − 1}. The curve shows explicitly over which range of powers the log-likelihood changes most drastically; when *β* is small and thus the importance distribution is close to the prior. Hence, a good numerical approximation of the log-likelihood curve is obtained when most powers take small values.

Draws from the power posterior distribution are obtained by running a modified Markov chain Monte Carlo (MCMC Metropolis *et al*., 1953; Hastings, 1970) algorithm:

1. Let *θ*_*j*_ denote the current parameter values at the *j*^*th*^ iteration, initialized at random at the start of the MCMC algorithm.
2. Propose a new values *θ′* drawn from a proposal kernel with density *q* (*θ′*|*θ*_*j*_).
3. The proposed state is accepted with probability

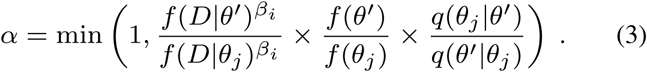
4. Set *θ*_*j*+1_ = *θ′* with probability *α* and to *θ*_*j*+1_ = *θ*_*j*_ otherwise.

As can be seen from this brief description of the modified MCMC algorithm, only the likelihood values need to be raised to the power *β*_*i*_. All remaining aspects of the MCMC algorithm stay the same as the standard implementations in Bayesian phylogenetics (Huelsenbeck and Ronquist, 2001; Drummond and Rambaut, 2007; Lakner *et al*., 2008; Lartillot *et al*., 2009; Höhna and Drummond, 2012).

It is important to note that every MCMC simulation for each power *β*_*j*_ ∈ *β* necessarily includes its own burn-in period before the first sample can be taken. The power posterior analysis can be ordered to start from the full posterior (*β*_*K*−1_ = 1.0) and then to use monotonically decreasing powers until the prior (*β*_0_ = 0.0) has been reached. Thus, the last sample of the previous power posterior run can be used as the new starting state. This strategy has been shown to be more efficient because it is easier to disperse from the (concentrated) posterior distribution to the (vague) prior distribution thereby reducing the burn-in period significantly (Baele *et al*., 2012).

### 2.2 Parallel power posterior analyses

The sequential algorithm of a power posterior analysis starts with a preburnin phase to converge to the posterior distribution. Then, consecutive power posterior simulations are performed sequentially, starting with *β*_K−1_ = 1.0 (*i.e.*, the posterior) to *β*_0_ = 0.0 (*i.e.*, the prior). Each power posterior simulation contains *L* iterations, with the likelihood of the current state recorded every *T*th iteration. These ‘thinned’ samples are less correlated than the original draws from the MCMC simulation. The number of samples taken per power is *n* = *L/T*. At the beginning of each run a short burn-in phase is conducted, for example 10% or 25% of the run length.

The parallel algorithm for a power posterior analysis is set up almost identically to the sequential algorithm (see Figure 2). Let us assume we have *M* CPUs available. Then, we split the set of powers into *M* consecutive blocks; the *m*^*th*^ block containing the powers from 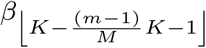 to *β*_⌊*K*−(*mK*/*M*)⌋_, *e.g.*, the first out of four blocks for 128 analyses contains {*β*_127_,…,*β*_96_}, the second block contains {*β*_95_,…,*β*_64_}, etc. If the set of *β* cannot be split evenly into blocks then some blocks have one additional simulation, which is enforced by using only the integer part of the index. This block-strategy ensures that each CPU works on a set of consecutive powers which has the advantage of a shorter burn-in between simulations because the importance distributions are more similar to one another.

**Fig. 2.**
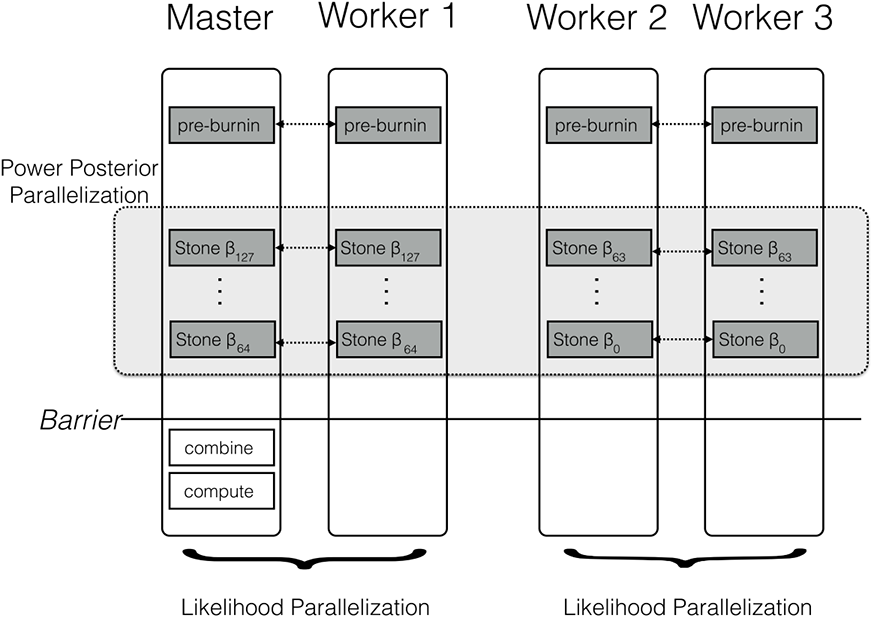
Schematic of the parallelization and workload balance between the master CPU and the worker CPUs. In this example we have *M* = 4 CPUs and *K* = 128 power posterior simulations (stones). The first CPU is the designated master and the remaining CPUs are the workers/helpers. The power posterior simulations are divided into two blocks from *β*_127_ to *β*_64_ and *β*_63_ to *β*_0_. The first two CPUs work on the first block of power posterior simulations and the last two CPUs work on the second block. Each pair of CPUs shares the likelihood computation between them. Each CPU starts with its own pre-burnin phase. Then, each CPU runs its block of power posterior simulations. Finally, the master combines the likelihood samples and computes the marginal likelihood estimate. Thus, the only barrier is after all the single power posterior simulations, which is after each single CPU has finished its respective job.

Regardless, each parallel sampler needs to start with an independent pre-burnin phase which creates an additional overhead. Thus, instead of running only one pre-burnin phase, as under the sequential power posterior analysis, we need to run *M* pre-burnin phases. This overhead could be removed only if it would be possible to draw initial values directly from the power posterior distribution.

Figure 2 shows a schematic of our parallelization algorithm. After the initial pre-burnin phase, the workload is divided into blocks and equally distributed over the available CPUs. Note that CPUs can be combined for distributed likelihood computation. No synchronization or communication between samplers is necessary because each power posterior simulation is independent. The only parallelization barrier occurs at the end when all power posterior simulations have finished. Finally, the master CPU collects all likelihood samples, combines the results, and computes the marginal likelihood using one of equations given below. These equations are computationally cheap compared with obtaining the likelihood samples. We thus expect that the performance gain is close to linear with the number of available cores. The algorithm described here is implemented in the open-source software `RevBayes` (Höhna *et al*., 2014; Höhna *et al*., 2016), available at http://www.RevBayes.com.

### 2.3 Path-Sampling

Path-sampling was the first numerical approximation method developed for marginal likelihood computation in Bayesian phylogenetic inference (Lartillot and Philippe, 2006). Path-sampling uses the trapezoidal rule to compute the integral of the log-likelihood samples between the prior and the posterior (see Figure 1), which equals the marginal likelihood (Lartillot and Philippe, 2006). The equation of the trapezoidal rule for a single likelihood sample from each power posterior simulation is

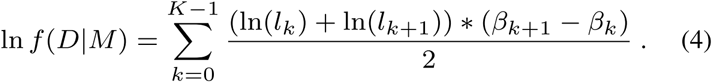

Samples of the log-likelihood have a large variance. Hence, it is more robust to take many log-likelihood samples and use the mean instead. This yields the equation to estimate the marginal log-likelihood,

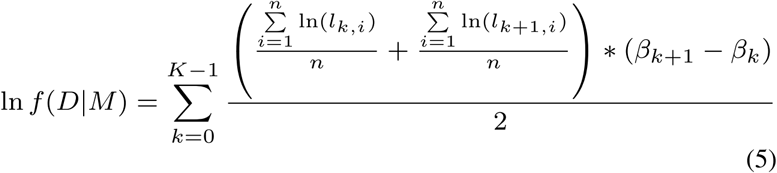

which was proposed by Baele *et al*. (2012).

### 2.4 Stepping-Stone-Sampling

Stepping-stone-sampling approximates the marginal likelihood by computing the ratio between the likelihood sampled from the posterior and the likelihood sampled from the prior. However, this ratio is unstable to compute and thus a series of intermediate ratios is computed: the stepping-stones (Xie *et al*., 2011; Fan *et al*., 2011). The stepping-stones can be chosen to be exactly the same powers as those used for path-sampling. The equation to approximate the marginal likelihood using stepping stone sampling is

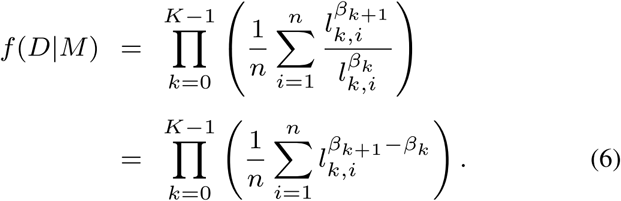

Numerical stability of the computed marginal likelihood can be improved by retrieving first the highest log-likelihood sample, denoted by max_*k*_, for the *k*^*th*^ power. Re-arranging Equation 6 accordingly yields

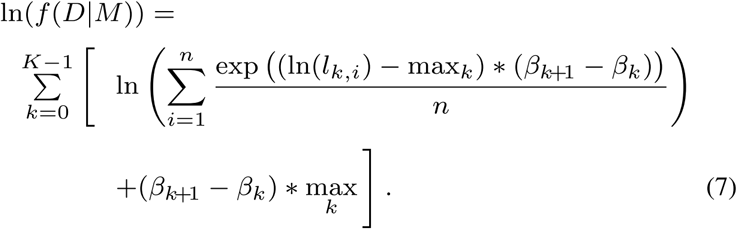

As seen in Equation 5 and Equation 7, only the set of likelihood, or log-likelihood, samples is needed to approximate the marginal likelihood. Both marginal likelihood estimates approach the true marginal likelihood when the number of samples and powers increases. Since both computations are comparably fast, they can be applied jointly and, for example, be used to test for accuracy without additional time requirements.

### 2.5 Simulation design

The objective of the simulation study was to test the performance gain when using multiple CPUs. Thus, we tested the performance of the parallel power posterior analyses using two phylogenetic examples; a smaller and a larger dataset. As the small example dataset we chose 23 primate species representing the majority of primate genera. We used only a single gene sequence, the cytochrome b subunit, containing 1141 base pairs. For the large example data set we chose an alignment with 4 genes from 305 taxa of the superfamily *Muroidea* (Schenk *et al*., 2013). For both examples we used the same model with the only difference that the larger dataset was partitioned into four subsets of sites. We assumed that molecular evolution can be modeled by a general time reversible (GTR) substitution process (Tavaré, 1986) with four gamma-distributed rate categories (Yang, 1994). Furthermore, we assumed a strict, global clock (Zuckerkandl and Pauling, 1962) and calibrated the age of the root. As a prior distribution on the tree we used a constant-rate birth-death process with diversified taxon sampling (Höhna *et al.*, 2011; Höhna, 2014) motivated by the fact that one representative species per genus was sampled, which is clearly a non-random sampling approach. Further details of the model can be found in the supplementary material.

Each analysis consisted of a set of *K* = 128 power posterior simulations (see Figure 2 for a schematic overview). The analyses started with a pre-burnin period of 10,000 iterations to converge to the posterior distribution. Then, each power posterior analysis was run for 10,000 iterations and samples of the likelihood were taken every 10 iterations. The 25% initial samples of each power posterior distribution were discarded as additional burnin. The marginal likelihood was estimated using both path-sampling and stepping-stone-sampling once all power posterior simulations had finished as they contribute to performance overhead in practice. We ran each analysis 10 times and measured the computation time on the San Diego Supercomputer (SDSC) Gordon. The experiment was executed using 1, 2, 4, 8, 16, 32 and 64 CPUs, respectively.

## 3 Results

We present the results of the average runtime as a function of the number of CPUs used in Figure 3. Performance gains are most pronounced when few CPUs are used. The runtime is almost halved when compared between 1 and 2 CPUs or 2 and 4 CPUs. For example, our primate analyses took on average 11.39 hours when using only a single CPU. By contrast, the analyses took only 5.95 hours and 3.15 hours when we used 2 CPUs and 4 CPUs respectively. Virtually the same runtime improvements were achieved for the larger *Murdoidea* dataset (Figure 3).

**Fig. 3.**
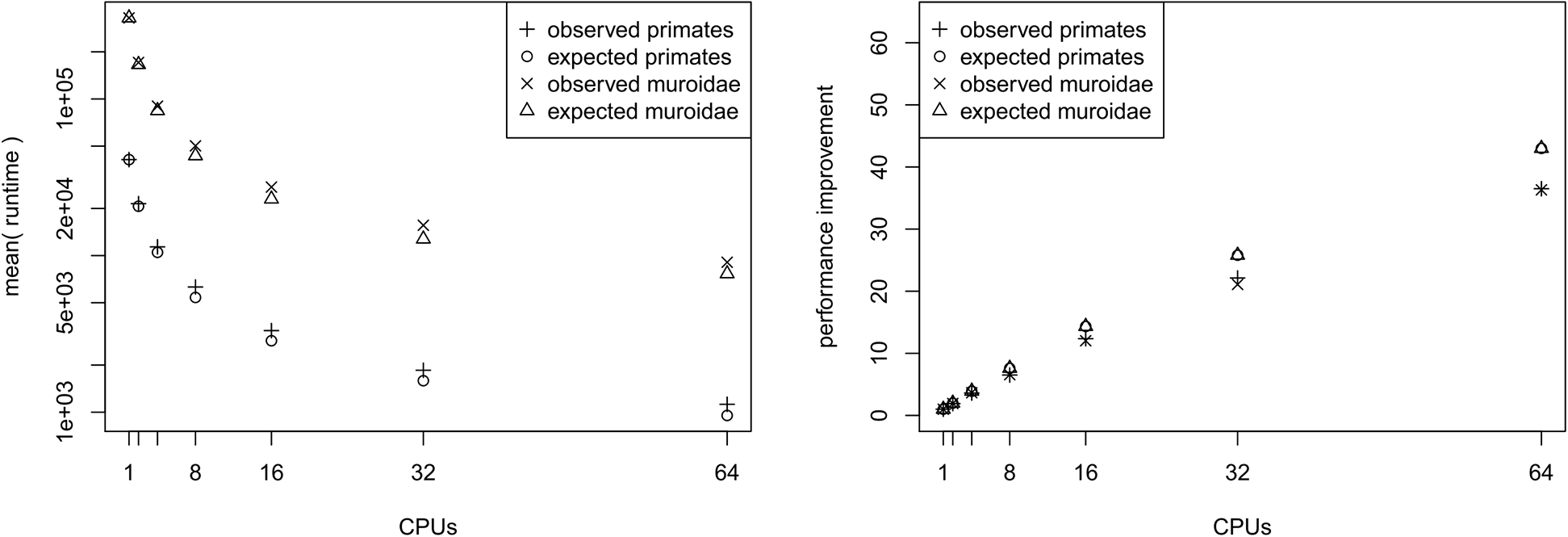
The average runtime of a marginal likelihood estimation on a simple phylogenetic model recorded over 10 repeated runs. The analyses were performed on the San Diego supercomputer cluster Gordon using 1, 2, 4, 8, 16, 32 and 64 CPUs. The runtimes were measured in seconds. The left graph shows the mean runtime as a function of the number of CPUs. The right graph shows the performance increase (fraction of time needed) compared with a single CPU. Both graphs show the actual performance increase and the expected performance increase (if there were no overhead between CPUs).

The performance increase levels off quickly once 8 or 16 CPUs are used. This is simply due to the fact that twice as many CPUs are needed each time to roughly halve the computational time. Hence, the gain from 1 to 4 CPUs is approximately equivalent to the gain from 16 to 64 CPUs. Additionally, the overhead (*i.e.*, the independently run pre-burnin for each chain) which each CPU needs to perform reduces the performance gain for larger number of CPUs.

We computed the expected runtime to assess whether our implementation achieved the largest possible performance gain. For example, we wanted to explore if there is an additional overhead for using parallelization that was possibly introduced by our specific implementation. Having *M* CPUs available, each CPU needs to run at most ⌈*K/M*⌉ power posterior simulations, which is the ratio of the total number of power posterior simulations to CPUs rounded upwards (ceiling). Additionally, each CPU runs its own pre-burnin phase, which had the same length as a single power posterior simulation in our tests. Therefore, we can compute the average runtime of a single power posterior simulation by dividing the runtime of the single CPU analysis by *K* + 1. Then, the expected runtime for *M* CPUs, *t*_*M*_, is given by

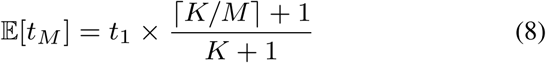

where *t*_1_ corresponds to the runtime when only one CPU was available. In general, our implementation seems to perform close to the expected optimal performance (Figure 3). However, we observe an increasing discrepancy between the expected and the observed performance gain when many CPUs were used. This discrepancy is most likely due to bottlenecks in competing hardware allocations. For example, we noticed that I/O operations performed on a network filesystem, which are commonly used among large computer clusters, significantly influenced the performance, especially when many CPUs frequently wrote samples of the parameters to a file.

We performed an additional performance analysis where we omitted the pre-burnin phase. This scenario could be realistic when one has already performed a full posterior probability estimation and only wants to compute the marginal likelihoods for model selection. In this case, the samples from the posterior distribution can be used to specify starting values of the power posterior analysis. Here we see that the performance improvement becomes more linear with the number of CPUs (see Figure 4). Although this case might not happen frequently in practice, we use this to demonstrate that only the pre-burnin phase prevents us from having an almost linear, and thus optimal, performance increase.

**Fig. 4.**
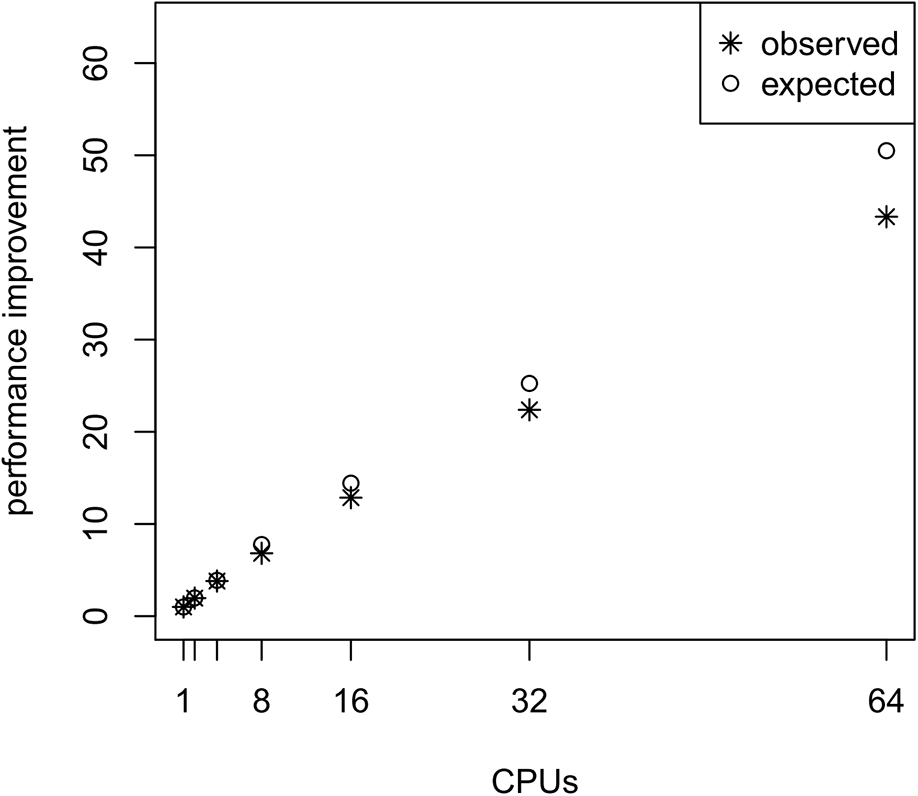
The average performance improvement (runtime reduction) when estimating the marginal likelihood on a simple phylogenetic model without pre-burnin phase, recorded over 10 repeated runs. The analyses were performed on the San Diego supercomputer cluster Gordon using 1, 2, 4, 8, 16, 32 and 64 CPUs. The runtimes were measured in seconds. The graph shows the actual and the expected performance increase compared with a single CPU, where performance is nearly linear

We also investigated whether the performance overhead (observed in Figure 3) is correlated with the number of stepping stones per CPU. For example, we observed the largest difference between the expected and actual runtime when 64 CPUs were used (each CPU ran only one or two power posterior simulations plus the pre-burnin phase). Thus, we tested if there was an effect of small numbers of power posterior simulations by running analysis with *K* ∈ {2, 3, 5, 10, 20, 30, 40, 50} on a single CPU. As the expected runtime, we computed the mean runtime per individual power posterior simulation when *K* = 50. Our results, shown in Figure 5, demonstrate that there is an intrinsic overhead for small number of power posterior simulations. This overhead seemed to be the cause of the discrepancy between our expected and observed performance increase in the parallel power posterior algorithm (Figure 3). Part of the overhead is caused by the additional time to start the process, load the data, allocate memory, receive file handles and all other tasks that need to be performed before and after a power posterior analysis.

**Fig. 5.**
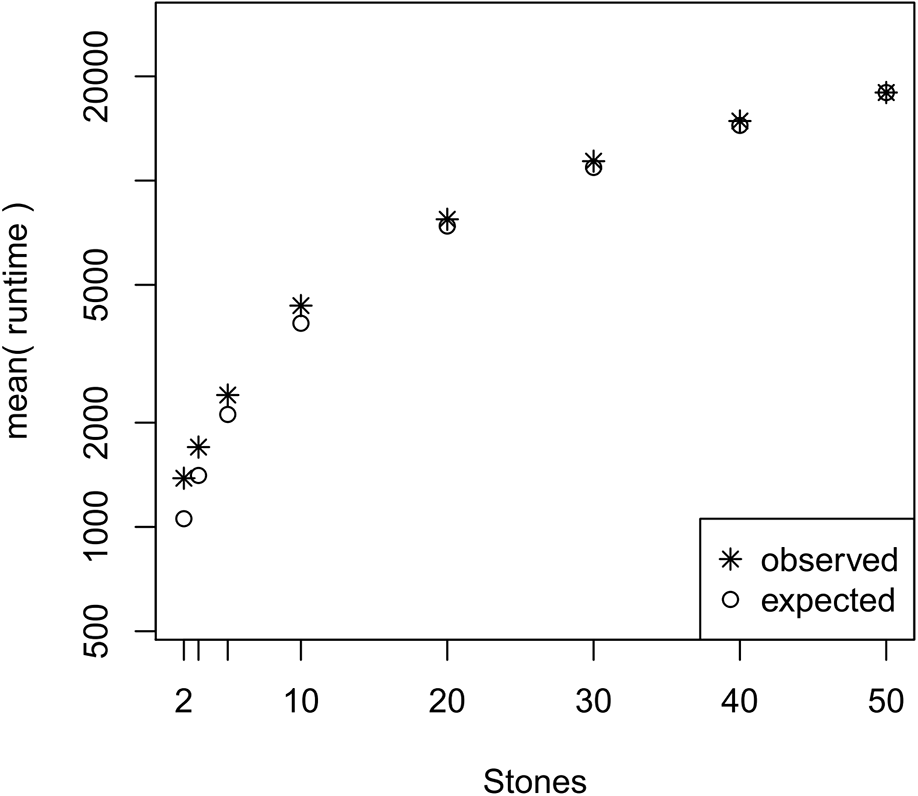
The average runtime over 10 repeated runs of a marginal likelihood estimation on a simple phylogenetic model for different number of powers posterior simulations *K*. The runtimes were measured in seconds. The graph shows the actual runtime and the expected runtime which is based on the mean runtime per power posterior simulation when *K* = 50.

Finally, we compared the performance increase when parallelizing the power posterior analysis, the likelihood computation, or both. For this combined parallelization scheme we implemented a hierarchical parallelization structure as describe by Aberer *et al*. (2014). For example, when 4 CPUs are available we can divide the likelihood computation over 2 CPUs and divide the power poster analysis into 2 blocks (see Figure 1). This test thus includes the parallelization approach over the likelihood function as suggested by Baele and Lemey (2013). We tested the performance difference using *M* = {2, 4, 8, 16, 32, 64} CPUs of which we assigned *N* to share the likelihood computation. We observed the best overall runtime reduction when we applied a combined likelihood and power posterior analysis parallelization (Table 1 and Table 2). Furthermore, the improvement of each parallelization yields diminishing returns when many CPUs are used, which additionally supports the utility of a combined parallelization scheme. We conclude that using 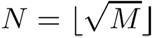 will give the overall best performance and set this distribution of CPUs as the default option in `RevBayes`.

**Table 1.** Runtime using *M* CPUs (rows) of which *N* CPUs (columns) are assigned to the likelihood computation. Here we show the results of the primates dataset.

| $M \setminus N$ | 1 | 2 | 4 | 8 | 16 | 32 | 64 |
| --- | --- | --- | --- | --- | --- | --- | --- |
| 2 | 21768 | 22268 | - | - | - | - | - |
| 4 | 11275 | 11434 | 12088 | - | - | - | - |
| 8 | 6253 | 6136 | 6185 | 6969 | - | - | - |
| 16 | 3336 | 3189 | 3162 | 3562 | 4612 | - | - |
| 32 | 1856 | 1709 | 1651 | 1846 | 2393 | 4738 | - |
| 64 | 1112 | 944 | 880 | 966 | 1217 | 2406 | 11363 |

**Table 2.** Runtime using *M* CPUs (rows) of which *N* CPUs (columns) are assigned to the likelihood computation. Here we show the results of the Muroidea dataset.

| $M \setminus N$ | 1 | 2 | 4 | 8 | 16 | 32 | 64 |
| --- | --- | --- | --- | --- | --- | --- | --- |
| 2 | 171797 | * | - | - | - | - | - |
| 4 | 92858 | 92212 | 90432 | - | - | - | - |
| 8 | 52329 | 51072 | 50244 | 53411 | - | - | - |
| 16 | 28248 | 26426 | 26792 | 27573 | 29297 | - | - |
| 32 | 15418 | 14423 | 14126 | 14599 | 15371 | 18450 | - |
| 64 | 9365 | 8234 | 7705 | 7649 | 8173 | 9641 | 18272 |

\* Runs using $M = 2$ CPUs with $N = 2$ CPUs per likelihood did not finish within the wall-time provided by XSEDE.

## 4 Conclusion

Modern phylogenetic analyses depend on increasingly complex models and increasingly large data set sizes. Even phylogenetic analyses which do not use molecular sequence data (for example, diversification rate analyses (FitzJohn, 2012), continuous trait analyses (Uyeda and Harmon, 2014), and historical biogeography analyses (Landis *et al*., 2013)) have grown more complex and use time-intensive likelihood calculations that are not always easily parallelizable. Both trends lead to longer runtimes, which is even more pronounced for Bayesian model selection exercises using marginal likelihoods; the path-sampling and stepping-stone-sampling algorithms used for approximating marginal likelihoods are inherently computationally demanding. In the present paper we have developed a simple parallel algorithm to speed up the computation of marginal likelihoods for Bayesian phylogenetic inference. In our simulation study, which serves mostly as a proof of concept, we showed that performance improvement is close to linear for few CPUs, *i.e.*, between one and 16 CPUs. An analysis that previously took 8 weeks on a single CPU can now be completed in four days when 16 CPUs are available.

Our new parallel power posterior analysis can be more than an order of magnitude faster than ordinary, sequential algorithms. The presented parallel algorithm should be straightforward to be implemented in other software or applied to a variety of different model types. Finally, the described parallelization scheme should be applicable to alternative methods for computing marginal likelihood (*e.g.*, Fan *et al*., 2011) and Bayes factors (Lartillot and Philippe, 2006; Baele *et al*., 2013) because all these approaches rely on a set of power posterior analyses.

## Acknowledgements

We would like to thank David Posada and three anonymous reviewers for valuable comments on a previous version of this manuscript.

